# Visual imagery vividness declines across the lifespan

**DOI:** 10.1101/2021.12.06.471435

**Authors:** Erzsébet Gulyás, Sára Sütöri, Andrea Lovas, Gergő Ziman, Ferenc Gombos, Ilona Kovács

## Abstract

The capacity to elicit vivid visual mental images varies within an extensive range across individuals between hyper- and aphantasia. It is not clear, however, whether imagery vividness is constant across the lifespan or changes during development and later in life. Without enforcing the constraints of strict experimental procedures and representativity across the entire population, our purpose was to take a first look at the self-reported level of imagery vividness and determine the relative proportion of aphantasic/hyperphantasic participants in different age-groups. Relying on the frequently used Vividness of Visual Mental Imagery questionnaire, we collected data on a random sample of 2252 participants between the ages of 12 to 60 years. We found a novel developmental pattern describing a declining ability to elicit vivid visual mental images in the group averages of different age-groups from adolescence to middle age. This effect involves both a decreasing proportion of individuals with very vivid imagery and an increasing proportion of individuals with weak imagery as maturation (assessed by boneage estimations in adolescents) and aging progresses. This finding may help to shed light on yet unknown developmental mechanisms of our internal, stimulus-independent processes, and might also help to determine genetic, maturational, and age-dependent factors in the extreme cases of hyper- and aphantasia.

## 1. Introduction

There has been a recent surge of both scientific and general interest in aphantasia. The term, coined by Zeman, Dewar, and Della Sala (Zeman et al., 2015), refers to a lack of mental visualization, or visual mental imagery, a capability readily available for most people. Two recent studies (Milton et al., 2021; Zeman et al., 2020) highlighted both extreme ends of the spectrum of mental imagery vividness: aphantasia on one end, and hyperphantasia – mental images as vivid as real seeing – on the other. This spectrum of mental imagery vividness is also reflected in Sir Francis Galton’s 1880 report, credited as the first scientific investigation of mental imagery (MacKisack et al., 2016), which gives an account of people with a complete lack of mental images, stretching to those with internal images “as clear as the actual scene” (Galton, 1880).

Researchers have been convinced since the 19th century that the vividness of mental visual imagery varies within an extensive range between individuals, and it also seems to be popular knowledge that the “internal eye” of a person might be very sharp in one case, and extremely weak in another. The question arises, however, whether imagery vividness is constant across the lifespan, or perhaps it changes during development and later in life. And this will be the topic of our explorations, driven by the inconsistencies in the current scientific literature on the issue.

Although suffering from validity issues, self-report surveys are a convenient way to assess the strength of the extremely subjective phenomenon of visual imagery, and many current studies rely on them. Perhaps the most widely used self-report questionnaire is the Vividness of Visual Imagery Questionnaire or VVIQ (Marks, 1973). In fact, the author of this popular questionnaire, David F. Marks, and his colleague, Anne R. Isaac attempted to explore vividness changes between 7 to 78 years of age using VVIQ (Isaac & Marks, 1994). They reported an increase of imagery vividness between 7 to 12 years of age, but no changes later in life. It is not clear, however, whether the early change seen in this study is due to the developing language comprehension abilities necessary to understand the entire complexity of the task or is it a true imagery related effect. Because of the low number of participants in each age-bin, the conclusion that imagery vividness is constant beyond 12 years of age is not very persuasive either. A preadolescent increase in imagery skills with a later permanence at this early-acquired level is also in contradiction with results on the less well-studied end of the spectrum of extreme hyperphantasia. It has been claimed that hyperphantasia is more frequent in elementary-school-aged children than in the older age groups (Haber & Haber, 1964; Giray et al., 1976; Haber, 1979). Additional studies attempting to see changes in mental imagery vividness across the lifespan seem to also suffer from the issue of small sample sizes, and thus, small power (Campos & Sueiro, 1993; Kemps & Newson, 2005; White et al., 1977; Wolmer et al., 1999). Some of these studies employed ranges spanning twenty years or more for the studied age-groups (Campos & Sueiro, 1993; White et al., 1977) which may conceal age-related changes on a finer scale. It seems, therefore, that the question with respect to lifespan changes of mental imagery is not very well answered by previous literature relying on questionnaires.

Given the increasing popularity of aphantasia and the growing number of papers in the past few years on this topic, it might be an interesting avenue to rely on existing data with respect to the prevalence of aphantasia in different age-groups. Equivalent proportions of aphantasic individuals across age-groups might either indicate that the condition is largely inherited while a potential change in prevalence should reveal developmental or aging effects, at least in the low vividness part of the spectrum. According to recent prevalence estimates, there are calculations from 0,7% (Milton et al., 2021; Zeman et al., 2020) to 2-3% (Faw, 2009), indicating a remarkable discrepancy between studies. While these numbers are related to absent mental imagery, the prevalence of people with imagery that is only dim/vague could reach 8,2% (Faw, 2009). The wide-ranging difference in the distribution of aphantasic individuals shows the lack of accordance about the precise terminology and could point out the shortage of representative samples. The dilemma of representativity also refers to the lack of research in the developmental stages. Most of the papers in the topic are engaged in undergraduate or adult participants (for recent examples, see: (Keogh & Pearson, 2014); Keogh et al., 2021; Milton et al., 2021; (Wicken et al., 2021), which sampling, however, ignores the potential effects in other age groups, including children and the elderly community. In aphantasia research, therefore, it is not easy to find even partial answers to our original question about imagery vividness across the lifespan.

In this study, we would like to explore the issue of lifespan changes in imagery vividness in a relatively large-sample study. Since this is an exploratory study, for feasibility reasons, our choice is to carry out a cross-sectional study instead of a longitudinal one. We will look at the potential changes in the proportion of people exhibiting different levels of imagery vividness across different age-groups from adolescence to middle age. Changes in the relative proportion of e.g., aphantasia might indicate a developmental or an aging effect. In this first exploratory large-sample lifespan study of imagery vividness, we will not include those age-groups that might require extra care because of the wide range of confounding factors such as the already mentioned language comprehension skills in children or the impact of medication in older age.

In terms of the methodological tools that can help address the question of age-related changes in imagery vividness, we are aware that there are relatively objective assessment options (Milton et al., 2021; J. Pearson et al., 2008, 2011), however, those entail tests of binocular vision in a laboratory environment and may not be easily adaptable to the changing developmental requirements (Ziman et al., 2021), not to mention feasibility issues on larger populations. Therefore, we opted for the most frequently used method to assess the vividness of internal images, and relied on the already mentioned VVIQ (Marks, 1973; McKelvie, 1995). Without enforcing the constraints of strict experimental procedures and representativity across the entire population, our purpose is to take a first look at the self-reported level of imagery vividness and determine the relative proportion of aphantasic/hyperphantasic participants in different age-groups. Our definition of aphantasia and hyperphantasia is very pragmatic, and it may not perfectly overlap with the usual connotations of these terms. We categorise participants by predetermining cutpoints on the VVIQ scale, and we simply look at the changes in the proportions of participants within these categories across age-groups. To our great surprise, there seems to be a very consistent decline in the vividness of mental imagery from adolescence through young adulthood and adulthood to middle age. We also look at vividness changes in terms of biological age in adolescents and find a maturational effect. These results might be an excellent starting point for further studies confirming our discovered age- and maturity-related imagery vividness effects and determining the neural background of those.

## 2. Materials and Methods

### 2.1. Participants

We recruited Hungarian participants aged 18 and above online. Participants filled out the VVIQ at the research group’s website (afantazia.hu). As we planned to compare results of at least four age-groups between 18 and 60 years of age, and analyse the data for the two sexes independently, we conducted a two-tailed power analysis (G*Power 3.1.9.7 software, Faul et al., 2007) to determine the minimum number of subjects required in each subgroup, aiming for at least a medium effect size (ES=0.5). We found that with an alpha of .05 and a power of 0.95, we need a minimum of 105 subjects in each subgroup that was achieved after receiving 2923 responses at the website. 557 responses had to be omitted due to missing gender and/or age information. 194 responses were omitted because participants indicated ages outside of our intended range. After the omissions, responses from 2172 participants remained, and we had at least 105 participants within each bin in the 21 to 60 years range, satisfying the above calculations (see Table 1.).

**Table 1.** Age ranges, descriptive statistics, and age distributions for the 5 age-groups.

| *age-group* | Age (years) |  |  |  | N |  |  |
| --- | --- | --- | --- | --- | --- | --- | --- |
|  | *min* | *max* | *mean* | *std.dev.* | *male* | *female* | *total* |
| 14 | 12 | 16 | 14.39 | 1.17 | 24 | 56 | 80 |
| 25 | 21 | 30 | 26.91 | 2.56 | 275 | 243 | 518 |
| 35 | 31 | 40 | 35.49 | 2.93 | 452 | 335 | 787 |
| 45 | 41 | 50 | 44.90 | 2.73 | 398 | 250 | 648 |
| 55 | 51 | 60 | 54.26 | 2.64 | 105 | 114 | 219 |

In addition to recruiting adult participants, we also invited 145 adolescents to fill out the VVIQ questionnaire online. These adolescents have gone through skeletal ultrasonic bone age assessment to determine their biological age (Kovács et al., 2021). Since the gender ratio of this sample was 1:2 (with twice as many girls than boys), we assumed this ratio in the power calculations, and aimed for an ES of 0.8 with a power of 0.9. 80 adolescents, between 12 to 16 years of age, responded to our invitation and filled out the questionnaire. This resulted in almost perfectly satisfying the minimal requirement of 25 boys and 51 girls (see Table 1.).

Altogether, 2252 participants filled out the VVIQ between the ages of 12 to 60 years, rendered into 5 age-groups (see Table 1. for detailed information on the age-groups).

Ethical approval was provided by the Ethical Committee of Pázmány Péter Catholic University (PPCU), Budapest, Hungary.

### 2.2. Questionnaire

We used the Hungarian version of the Vividness of Visual Imagery Questionnaire (Marks, 1973; McKelvie, 1995) translated by our research group. VVIQ asks the participant to imagine a relative or friend, a sunrise, a shop, and a country scene. For each scenario, four details are given and asked to be incorporated in the visual mental image and evaluated according to their vividness afterward. Compared to the original version (Marks, 1973), we used a reversed scale (see also in Zeman et. al, 2015, 2020), where 1 indicates “*No image at all, you only “know’’ that you are thinking of the object*”, and 5 means “*Perfectly clear and lively as real seeing*”.

### 2.3. Bone-age assessment

Ultrasonic bone age estimation carried out by scanning the wrist area is a recently introduced measure of maturity in developmental research (Kovács et al., 2021), and provides a more reliable and accurate assessment of biological age than the previously used ones. We assessed skeletal maturity (bone age) with an ultrasonic device (Sunlight BonAge, Sunlight Medical Ltd, Tel Aviv, Israel). Ultrasonic bone age estimation was carried out at the school of the participants or at the Research Centre for Sport Physiology at the University of Physical Education, Budapest. The procedure is fully described in the paper by Kovács et al. (2021).

### 2.4. General procedure

A popular article has been published to disseminate ideas about aphantasia at an online Hungarian news portal (Molnár, 2020). The article included a link to the Hungarian version of the VVIQ questionnaire at the research website (afantazia.hu). Prior to accessing and filling in the questionnaire, volunteers had to provide written informed consent indicating that they are 18+ years old, understood the nature of the study and will receive no compensation afterwards.

An alternative approach was taken to involve adolescents with the aim to ensure necessary information gets to parents, who can then grant informed consent. For this study, underaged participants and their parents were contacted via email which included a link pointing to the same introductory questions followed by the Hungarian version of the VVIQ.

After consenting, participants received two introductory questions of the same nature, intending to foster understanding of the VVIQ questionnaire afterwards. First, we asked participants to imagine an apple, then a horse and indicate the vividness of the images in both cases. Participants found these two questions helpful to foster their understanding of what ‘seeing’ an image in one’s mind might mean. Following the introductory tasks, participants progressed to the Hungarian VVIQ and finally gave answers to demographic questions, such as age and gender.

### 2.5. Statistical analysis

We used one-way analysis of variance (ANOVA) to look for significant differences between age-groups and genders. We tested associations between VVIQ score and bone age by the Pearson product-moment correlation coefficients (TIBCO STATISTICA 13.5.0.17 software, TIBCO Software Inc., 2018). To find the best fitting curve for the declining age-group averages we used Microsoft Excel’s power curve fitting method. We calculated standard error of the mean for the VVIQ scores of the age-groups and genders. We calculated confidence intervals for the population proportions of aphantasic participants and of participants within each age-group providing VVIQ scores less than 20, between 20 and 40, between 40 and 60, and above 60 VVIQ score-range. Standard error of the mean and confidence intervals for the population proportions were calculated using Microsoft Excel.

### 2.6. Data availability

The analyzed dataset and supplementary files were made available at the datadryad.org repository.

## 3. Results

### 3.1. Developmental hyper- and aphantasia

Average VVIQ scores are shown for each age-group in Fig. 1.A. Imagery vividness, as represented by VVIQ scores, significantly drops every ten years between 14 and 45 years of age (14 vs. 25-year-olds: F = 34.86, p < 0.001; 25 vs. 35-year-olds: F = 13.39, p < 0.001; 35 vs. 45-year-olds: F = 17.79, p < 0.001). A power trendline almost perfectly fits the declining imagery vividness scores (R^2^ = 0.9948).

**Fig. 1.**
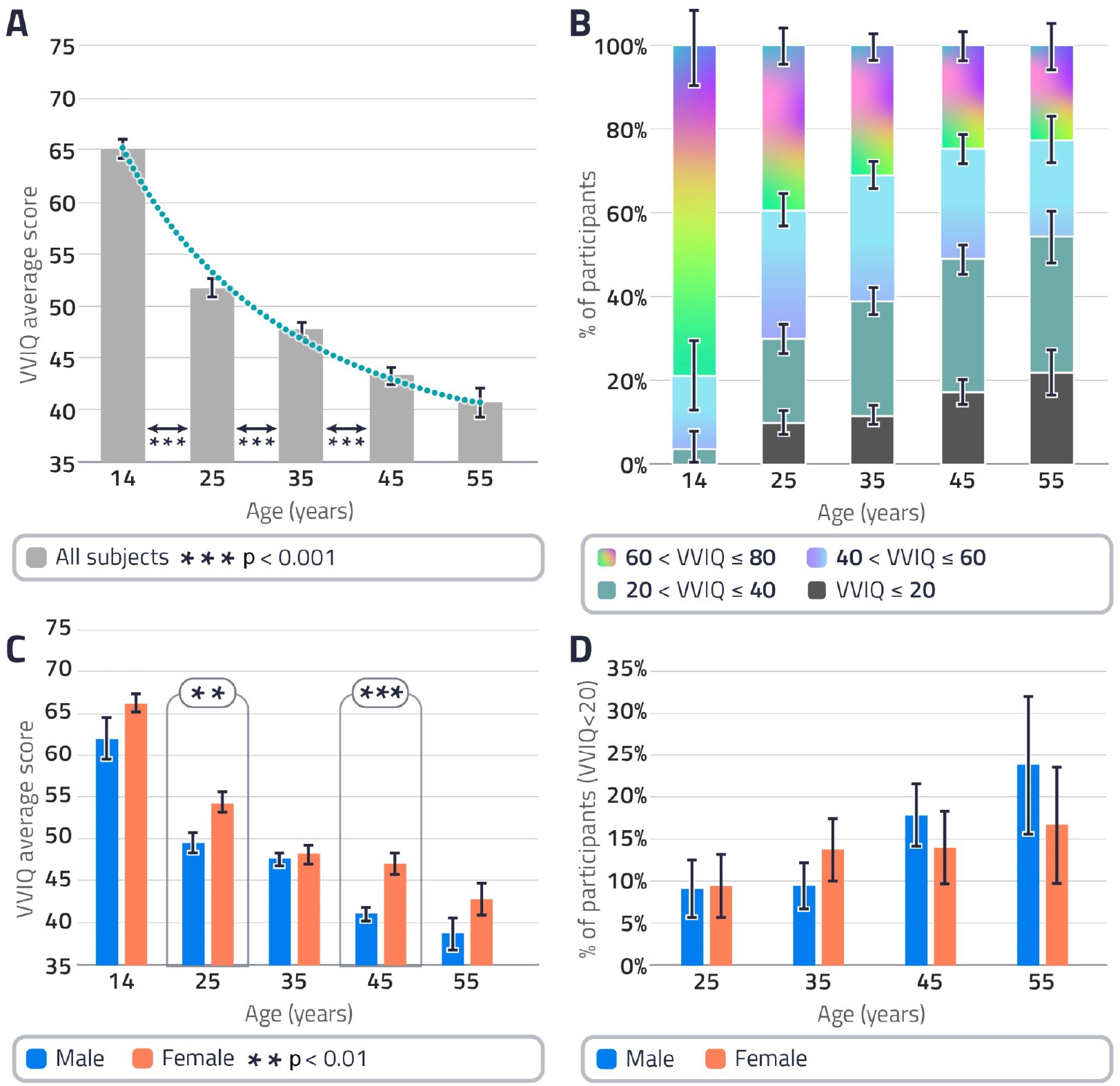
A. Grey columns represent average VVIQ scores within age-groups, based on the scores of each participant within an age-group, including aphantasic individuals (error bars are ±SE based on the ANOVA). The declining tendency of imagery vividness is highlighted with a significant power trend line (green dots, y = 64.434x^-0.285^, R^2^ = 0.9948). Vividness significantly drops every ten years until the age of 45. B. Coloured bars show the proportion of participants within each age-group providing VVIQ scores less than 20 (dark grey), between 20 and 40 (green), between 40 and 60 (turquoise), and above 60 (rainbow colours) VVIQ score-range. High scores indicate high imagery vividness. Bars indicate 95% confidence intervals for each range of scores, respectively. Participants in the rainbow-colour-range might be called self-reported hyperphantasic, while those in the dark-grey-range are likely aphantasic. Hyperphantasia seems to decline with age, while the proportion of aphantasic participants increases with age. C. Coloured columns show average VVIQ scores for the two sexes within each age-group, error bars are ±SE). Imagery vividness seems to decline faster in males, with a significant difference between males and females at the age of 25 and 45. D. Coloured columns show the proportion of aphantasic participants (VVIQ score lower than 20) for the two sexes (14-year-olds are not shown since there was no aphantasic participant in this age-group. Bars indicate 95% confidence intervals.

To see what might be behind the decline of the average score, i.e., whether it is because the proportion of people with vivid imagery is decreasing or perhaps because the proportion of aphantasic individuals is increasing, we divided the 80 points range of the VVIQ into four 20-point ranges (see Fig.1.B). Participants in the 60 to 80 points range might be called self-reported hyperphantasic, while those below 20 points are likely aphantasic. Since the minimal score on the VVIQ is 16 (1 point is given on each 16 items, which is the minimal point), the aphantasic range is narrower, and it is close to the extreme aphantasic range. However, we reasoned that it would be useful to keep it narrow to see the proportion of extreme aphantasics in each age-group and compare it to already existing data in the recent literature.

Fig. 1.B. reveals the distribution of participants within the four ranges of imagery vividness. 14-year-olds seem to stand out as 80% of this participant pool self-reported very high imagery vividness (scoring above 60) that we will call hyperphantasia. Hyperphantasia decreases to nearly 20% in 55-year-olds. As for the remaining age-groups, it seems that confidence intervals are not overlapping up to 45 years of age, indicating a sharply declining proportion of hyperphantasics between 14 to 45 years of age. There are no self-reported extreme aphantasics (scoring below 20) in the 14-year-old group, while more than 20% reports extreme low vividness in the 55-year-old group, as it is shown Fig.1. B. Based on the confidence intervals it seems that the most pronounced changes are occurring in late adolescence, and between 35 to 45 years of age in this respect. To conclude, while hyperphantasia is obviously on the decline with age, the proportion of aphantasic individuals also seems to increase with age.

The comparison of average VVIQ scores between females and males is shown in Fig.1. C. In general, females tend to have higher self-reported imagery vividness scores, while age-related vividness deterioration seems to be faster in males. The differences between the two sexes are significant in the 25 and in the 45-year-old groups (F = 7.55, p < 0.01; F = 14.49, p < 0.001).

Fig. 1.D. reveals the proportion of aphantasic individuals (scoring below 20) within each age-group, for the two sexes separately. Although the proportion of aphantasic individuals seems to be higher in males at older ages, there is no significant difference between males and females

### 3.2. Adolescent maturity and imagery vividness

As mentioned in the Methods section, our adolescent participants went through a bone age estimation procedure. Since bone age reflects biological maturity that not only determines bodily changes around puberty but is also responsible for cortical maturation, we analysed VVIQ responses as a function of bone age. The impact of biological maturity seems to be relevant only within a narrower temporal window, not across the entire adolescent chronological age-range. For the 33 participants with chronological age of 13 (N = 20) and 14 (N = 13) years, bone age had a significant impact on VVIQ scores, with more mature individuals providing lower scores. In other words, there is a negative correlation between bone age and VVIQ scores (r = -.36, p = .04, two-tailed; mean VVIQ score: 64.78, mean bone age: 14.16). There was no significant bone age effect in the other adolescent age-groups.

## 4. Discussion and conclusions

With respect to our original question whether imagery vividness is constant across the lifespan or changes during development and later in life, we found a surprising and yet undocumented declining tendency with age that is more pronounced in males. Highly vivid imagery that might be characterised as hyperphantasia is common during the teenage years, while the proportion of hyperphantasics is sharply declining from adolescence to middle age. Extreme aphantasia, non-existent in adolescents, seems to become increasingly prevalent in the later years. Additionally, in adolescents, advanced biological maturity is correlated with weaker imagery vividness. We interpret these findings as evidence for the waning of imagery vividness as a function of chronological age between adolescence and middle age, and as a function of biological age in adolescents.

We believe that the discovered developmental changes in imagery vividness in general, and in the prevalence of aphantasia, are novel findings. In terms of mental imagery and aging, there exist different experimental approaches, as well as studies mainly around mental rotation, visuo-spatial internal representation, and visual working memory (for recent examples, see:(Craik & Dirkx, 1992; Dror & Kosslyn, 1994; Isaac & Marks, 1994; Wimmer et al., 2015). However, none of these are engaged in developmental aspects of visual mental imagery vividness on its own. Earlier studies, mentioned in the introduction, using VVIQ in different age-groups (e.g.,Isaac & Marks, 1994; Campos & Sueiro, 1993; Kemps & Newson, 2005; White et al., 1977; Wolmer et al., 1999) suffer from the issues of small sample size, small power, poor temporal resolution on the age-scale, and other artefacts leading to inconsistent and uninterpretable findings. The above described, age-related declining pattern of imagery vividness seems to provide a fresh and more coherent picture, and perhaps an alternative way of thinking about the background of internal representations. The decreasing vividness may reveal a kind of intriguing neural plasticity in the context of the changing nature of mental representations during ontogeny. In line with this assumption, our observations could open new directions in the literature to further understand the neural correlates behind internal representations. Our results may also shed light on the necessity of more representative surveys in this field to get more persuasive information about the maturational effects of the phenomenon.

Our observation on the remarkable distribution of the two extremes – namely hyper- and aphantasia – is unique in the literature. About the phenomenon of aphantasia, most of the studies envisage aspects of its role in different kind of cognitive functions, and potential impairments or compensative internal processes, mainly only in adult samples (for recent examples, see: Jacobs et al., 2018; Pounder et al., 2018; Keogh et al., 2021; Milton et al., 2021; Wicken et al., 2021). However, to understand the phenomenon more comprehensively, it would be essential to examine the nature of lifelong prevalence as well. The fact that both chronological and biological age are negatively correlated with imagery vividness seems to uncover a developmental process and indicates the existence of “developmental aphantasia” in addition to the potentially genetically based and acquired forms.

The other terminus of the imagery vividness spectrum is the phenomenon of “extreme hyperphantasia” (Zeman, 2020). In contrast to aphantasia, this kind of extreme, ‘offline-perceptual’ behaviour means to have abnormally strong, or photo-like visual imagery. According to the previous prevalence calculations, this mental representational ability is more frequent in the group of elementary-school-aged children, than among subjects in other age groups (Giray et al., 1976; Haber, 1979; Haber & Haber, 1964). Different theoretical approaches exist to explain this distributional pattern of which the most popular viewpoint is the developmental hypothesis. According to this assumption, extreme imagery vividness is an early capacity, modulated or lost by the progression of developmental processes during childhood. Nevertheless, this aspect could not necessarily provide a reliable explanation to any longitudinal observations, namely to the cases within the eidetic subjects that remained eidetically classified in the entire experimental time interval (Leask et al., 1969). Based on our findings, we would like to suggest that in addition to the genetically based neurodiversity along the imagery spectrum, a lifelong developmental pattern of decreasing imagery vividness is also part of the picture. The mechanism behind this decline and the neural background is not within the scope of our investigation here, however, we hope to facilitate the more detailed investigation of those.

As it is illustrated in Figure 2., inspired by the current findings, we would like to propose that the vividness of visual mental imagery is shaped by developmental factors, and there is a natural tendency for less vivid mental images with both maturation and aging. Although we would like to leave the possibility open that there might be a small proportion of individuals with genetically based extreme hyper- or aphantasia whose imaging capacities are unaffected by maturational or developmental factors, we also claim the relevance of the developmental changes. As for future studies, this relevance might be twofold. On one hand, by acknowledging the changing distributions of imaging capacities, future studies might involve samples more representative for age and gender to determine the actual prevalence of either, potentially genetically based extremes. On the other hand, the clear declining tendency raises several questions about the developmental mechanisms that bring about such a change. Does it have any relevance in acquiring our human nature during development? Is it a necessary change coupled with our unique language learning abilities? Is there a difference between genetically based and developmental hyper- and aphantasia?

**Fig. 2.**
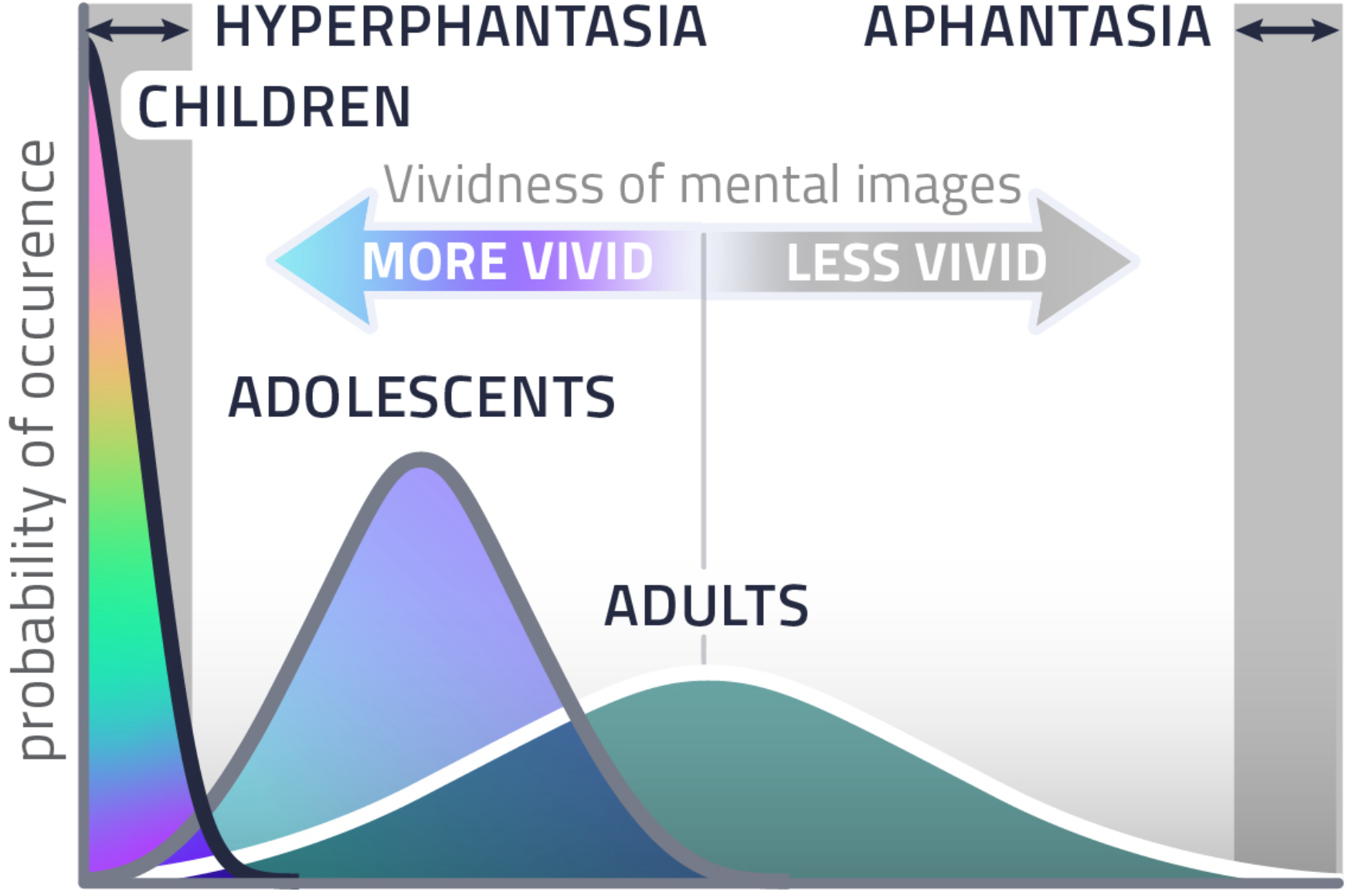
Predicted developmental trend for imagery vividness. While children are expected to fall within the ‘hyperphantasic’ range, adults are predicted to have less vivid imagery both on average, and in terms of absolute vividness. Adolescents should fall in between. The prevalence of ‘developmental aphantasia’ is expected to increase with age.

Fig. 2. Predicted developmental trend for imagery vividness. While children are expected to fall within the ‘hyperphantasic’ range, adults are predicted to have less vivid imagery both on average, and in terms of absolute vividness. Adolescents should fall in between. The prevalence of ‘developmental aphantasia’ is expected to increase with age.

Instead of indulging in further exciting but unanswered questions, let us also note the limitations of our exploratory study that could be overcome in further investigations. First, despite the large number of participants in the adult age-groups, our study cannot be considered representative, and it is not longitudinal. Therefore, we cannot completely rule out confounds related to random samples, and confounds that may include social, educational, technological, or lifestyle changes over time that may affect the spectrum of mental imagery vividness across the examined age groups. Attrition bias might also be present if, e.g., mortality would decrease in the presence of aphantasia, or increase in the general and hyperphantasic population, maybe due to a confounder with psychiatric or neurodegenerative diseases reducing life expectancy (Ji et al., 2019; D. G. Pearson et al., 2013; J. Pearson et al., 2015). It is also a shortcoming of this exploratory study that we used a self-report questionnaire that is on the one hand, subject to several response biases, and on the other hand, it may not be readily applicable in children. Since the extension of our original question to childhood seems extremely relevant, more objective measures involving lower levels of cognitive complexity in the responses are called for. For example, a no-report version (Ziman et al., 2021) of the binocular rivalry dominance priming method (Milton et al., 2021; J. Pearson et al., 2008, 2011) might be a useful paradigm in forthcoming studies.

To sum up, we found a novel developmental pattern describing a declining ability to elicit vivid visual mental images in the group averages of different age-groups from adolescence to middle age. It seems that this effect involves both a decreasing proportion of individuals with very vivid imagery and an increasing proportion of individuals with weak imagery as maturation and aging proceeds. We believe that this finding deserves further investigation as it may help to shed light on yet unknown developmental mechanisms of our internal, stimulus-independent processes, and might also help to determine genetic, maturational, and age-dependent factors in the extreme cases of hyper and aphantasia.

## Acknowledgements

We acknowledge support from NKFI (Nemzeti Kutatási, Fejlesztési és Innovációs Hivatal) 134370 to IK.

